# Rapid, direct, and sequence-specific identification of RNA viruses in various crop plants using CRISPR/Cas13a

**DOI:** 10.1101/2024.02.22.581525

**Authors:** Hagit Hak, Steffen Ostendorp, Anton Reza, Shany Ishgur Greenberg, Gur Pines, Julia Kehr, Ziv Spiegelman

## Abstract

Plant viruses are destructive pathogens causing significant damage to various crop species. Rapid, sensitive, and specific detection is crucial for the effective containment of emerging and resistance-breaking viruses. CRISPR/Cas has been established as a useful tool for plant virus identification. However, its application for on-site, direct detection of viruses from plant tissues is still limited. In this study, we present a rapid method for detecting viruses directly from RNA of different crop species using CRISPR/Cas13a. We successfully applied this method to identify tomato brown rugose fruit virus (ToBRFV) in infected tomato plants and differentiate it from closely related tobamoviruses. ToBRFV could be identified in a 100-fold dilution and early during infection, prior to the onset of viral symptoms. Moreover, CRISPR/Cas13a was used to directly identify cucumber green mottle mosaic virus (CGMMV) in cucumber plants and turnip mosaic virus (TuMV) in *Brassica napus* plants. Finally, we developed a user-friendly, extraction-free, 15-minute protocol for on-site ToBRFV identification using a portable fluorescent viewer and a mobile phone camera. This protocol was successfully applied for ToBRFV detection in a commercial greenhouse. These results demonstrate that CRISPR/Cas13a is a robust technology for direct, rapid, sensitive, and specific identification of multiple viruses in different crop plants that can be easily implemented for on-site detection.

## Introduction

Plant viruses are major agricultural pathogens that cause severe damage to various crop species, with an estimated global impact of 30 billion dollars (Jones and Naidu, 2019). The recent emergence of several new and resistance-breaking viruses poses a major challenge to global agriculture (Jones, 2021; Ristaino *et al*., 2021; Salem *et al*., 2023). Viruses of the *Tobamovirus* genus, including the tobacco mosaic virus (TMV) and tomato mosaic virus (ToMV) inflict destructive diseases that result in significant loss of yield and deterioration of fruit quality in different crop plants (Salem *et al*., 2023; Spiegelman and Dinesh-Kumar, 2023). Tobamoviruses are extremely infectious viruses transmitted by mechanical contact with working hands, agricultural tools, and soil. Recently, a new and emerging tobamovirus named tomato brown rugose fruit virus (ToBRFV) has caused significant damage to the global tomato industry. First originating in the Middle East, ToBRFV has now become a global pandemic, which severely affects the international commerce of tomato seeds, plants, and fruits (Salem *et al*., 2023; Zhang *et al*., 2022). Importantly, ToBRFV can be transmitted on the surfaces of seeds (Davino *et al*., 2020) and distributed by marketed tomato fresh fruit (Klap *et al*., 2020; Skelton *et al*., 2023) and pollinators (Levitzky *et al*., 2019; Skelton *et al*., 2023). Currently, ToBRFV is managed by strict regulation at key points of commerce, such as international borders, which require sensitive detection and monitoring of the virus (EPPO, 2021). Since ToBRFV symptoms are inherently similar to other tobamoviruses, such as TMV and ToMV, it is crucial for ToBRFV detection protocols to be highly specific and enable its distinction from related viruses. Therefore, the development of an accessible, rapid, sensitive, and species-specific method to identify ToBRFV is of great significance to the global tomato industry.

Another important member of the tobamovirus genus is the cucumber green mottle mosaic virus (CGMMV). This virus infects multiple crop species of the *Cucurbitaceae* family, including cucumber, squash, zucchini, and melon. While CGMMV has been characterized almost 90 years ago (Ainsworth, 1935), in recent decades, the dispersal of CGMMV has dramatically increased, and currently it poses a substantial threat to the global cucurbit industry (Dombrovsky *et al*., 2017). Since there is no commercially available genetic resistance to CGMMV, phytosanitary measures combined with frequent and sensitive diagnostics are required to control this virus in a manner similar to ToBRFV (Darzi *et al*., 2020).

An additional destructive group of plant viruses are members of the *Potyvirus* genus. Among potyviruses, turnip mosaic virus (TuMV) has the broadest host range, infects both monocotyledonous and dicotyledonous plants, and is transmitted by 79 species of aphids (Nellist *et al*., 2022). In *Brassica napus* (rapeseed; canola), TuMV can cause losses of up to 70% in yield (Hardwick *et al*., 1994). While several TuMV resistance genes have been characterized in *B. napus*, the recent emergence of resistance-breaking TuMV isolates (Guerret *et al*., 2017) may pose a significant threat to the canola oil industry.

The lack of genetic resistance against plant viruses in various crops requires strict management strategies that include a combination of early detection and phytosanitary measures. Therefore, simple and robust approaches for early, specific, and sensitive detection are required (Mehetre *et al*., 2021; Rubio *et al*., 2020). Currently, the most common technologies for plant virus identification rely on sequence-specific molecular tools, such as reverse transcription quantitative PCR (RT-PCR), digital droplet PCR (ddPCR), and high throughput sequencing (HTS). These technologies are highly sensitive and sequence-specific, however, their application is restricted to laboratories with specialized equipment and trained personnel (Mehetre *et al*., 2021; Rubio *et al*., 2020). Other, more robust tools are antibody-based approaches, such as enzyme-linked immunosorbent assay (ELISA) and rapid lateral flow detection kits. These technologies, which are based on polyclonal antibodies against the viral coat proteins, are more accessible, and some of them are field deployable. However, they are less sensitive than nucleic-acid-based technologies (Bhat *et al*., 2022). In addition, antibody-based techniques are less specific due to their cross-reactivity, which stems from the high conservation of viral proteins in closely related virus species. Therefore, there is an essential need to develop a direct, rapid, sensitive, and sequence-specific protocol for field-deployable identification of plant viruses.

One emerging technology for viral diagnostics is using Clustered Regularly Interspaced Short Palindromic Repeats (CRISPR) (Chertow, 2018; Yin *et al*., 2021). This technology relies on one of two CRISPR-associated (Cas) proteins, Cas12 or Cas13 for the detection of DNA and RNA, respectively. Cas enzymes are directed to their specific nucleic acid targets by forming a complex with CRISPR RNA (crRNA) molecules, complementary to the targeted sequence. Once Cas12 or Cas13 bind to their targets, they are activated, and in addition to the sequence-specific digestion, a non-specific collateral cleavage of single-stranded (ss)DNA or ssRNA molecules, respectively, is also achieved (Freije and Sabeti, 2021). This non-specific cleavage activity can be utilized for robust detection of nucleic acid sequences by labeling a ssDNA or ssRNA probe with fluorophore and quencher. Cas-mediated degradation of these molecules releases the fluorophore, thereby emitting a fluorescent signal. In human medicine, this technology has been proven effective for the diagnostics of multiple viral diseases (Chen *et al*., 2018; Gootenberg *et al*., 2017; Kaminski *et al*., 2021; Myhrvold *et al*., 2018). The emergence of the COVID-19 pandemic has boosted the development of CRISPR-based viral diagnostics tools, resulting in several commercial platforms for the rapid detection of SARS-CoV-2 (Rahimi *et al*., 2021; Safari *et al*., 2021). CRISPR/Cas12 technology has been demonstrated for detecting several plant viruses (Aman *et al*., 2020; Mahas *et al*., 2021; Marqués *et al*., 2022; Xu *et al*., 2023). Previously, we have established that Cas12 technology can be utilized for the specific detection of ToBRFV, distinguishing it from the closely related ToMV, and identifying it from samples collected in a commercial greenhouse (Alon *et al*., 2021). In a different study, this technology was paired with RT-loop-mediated amplification (LAMP) and lateral flow strips, demonstrating the potential of this technology for on-site diagnostics (Bernabé-Orts *et al*., 2022). It is important to note that all Cas12-based detection protocols for RNA viruses require reverse-transcription followed by amplification, which extends the time and complexity of these methods (Bernabé-Orts *et al*., 2022).

While Cas12 has been well-established for plant virus detection, the ability of Cas13 for plant virus diagnostics has been less explored. The ability of Cas13 to directly bind to specific RNA molecules suggests its potential for direct, amplification-free detection of plant RNA viruses. Previously, Cas13a has been shown to enable the detection of TSWV, however, this technology still requires amplification of the viral sequence (Zhang *et al*., 2021). A recent study showed that Cas13a could be utilized for the direct detection of RNA viruses in model plants, suggesting the potential of this system for rapid, direct, and field-deployable detection of viruses (Marqués *et al*., 2022).

Here, we present a rapid and robust approach for the detection of agriculturally-relevant RNA viruses in important crop species based on CRISPR/Cas13a technology. We have developed a direct, amplification-free, and species-specific method to identify ToBRFV in tomato plants. In addition, the protocol was used for the detection of CGMMV in cucumber plants and TuMV in *B. napus* plants. Finally, this method has been simplified into a 15-minute-long protocol for on-site detection of ToBRFV (Fig. 1) based on only two steps (1) Rapid RNA sampling from the infected plant (5 minutes) and (2) Incubation with the Cas13a reaction mixture (5-10 minutes). Samples can then be monitored using a simple, portable fluorescence viewer (Fig. 1). These results may pave the way for the development of a new generation of kits for field-deployable detection of various plant viruses.

**Figure 1.**
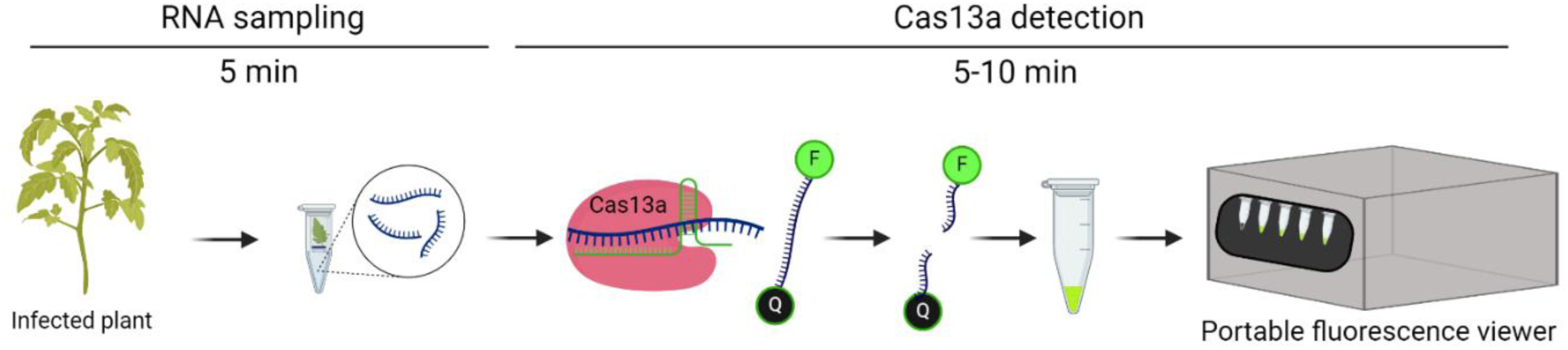
Rapid and direct diagnostics of RNA crop viruses using CRISPR/Cas13a. An illustration of crop virus detection using CRISPR/Cas13. RNA is sampled from an infected plant into a lysis buffer. The RNA is then directly incubated for 5-10 minutes with the reaction mixture which includes Cas13a, specific crRNA and an ssRNA reporter molecule, which consists of a fluorophore (F) and a quencher (Q). Once the viral RNA is detected, the RNA reporter is degraded the fluorophore is separated from the quencher, and a fluorescence signal is emitted. This signal can be viewed using a simple, portable, fluorescence viewer. The figure was created using Biorender.com.

## Results

### Species-specific detection of ToBRFV in tomato plants using CRISPR/Cas13a

Genomic RNA of tobamoviruses accumulates to very high concentrations in plant tissues (Supplementary figure S1). Based on this observation, we hypothesized that CRISPR/Cas13a could be used for amplification-free detection of ToBRFV in plants. To test the potential of Cas13a for direct and specific detection of ToBRFV, we designed three different crRNAs, targeting different regions in the ToBRFV genome (Figure 2a): *TB-crRNA1* targeted ORF1/2, encoding the viral replication protein(s), *TB-crRNA2* targeted ORF3, encoding the viral movement protein crRNA and *TB-crRNA3* targeted ORF4, which encodes the coat protein (Figure 2a). To allow species-specific detection, the crRNAs were designed to target sequences that are altered in ToBRFV compared to ToMV and TMV (Figure 2b). The three tested crRNAs were then incubated with RNA extracted from leaves of tomato plants infected with ToBRFV or ToMV, *Nicotiana benthamiana* plants infected with TMV, and healthy tomato plants (Figure 2c-e). These crRNAs were variable in their detection capacity: *TB-crRNA1* emitted a strong fluorescence signal with RNA from ToBRFV-infected plants, while no signal was observed in RNA from healthy, ToMV- or TMV-infected plants (Figure 2c), *TB-crRNA2* could not distinguish between ToBRFV- and ToMV infected plants (Figure 2d), and *TB-crRNA3* was able to specifically detect ToBRFV, however with reduced signal intensity as compared to *TB-crRNA1* (Figure 2e). In addition, we developed a crRNA for the specific detection of ToMV, which was able to distinguish it from ToBRFV- and TMV-infected plants (Supplementary figure S2). We further tested whether the combination of different crRNAs can improve ToBRFV detection capacity (Supplementary figure S3). The combination of *TB-crRNA1* and *TB-crRNA2* resulted in reduced detection as compared to *TB-crRNA1*. However, the combination of *TB-crRNA1* and *TB-crRNA2* generated enhanced detection as compared to *TB-crRNA1*. These results suggest that combining different crRNAs directed against the same virus can result in synergistic detection capacity.

**Figure 2.**
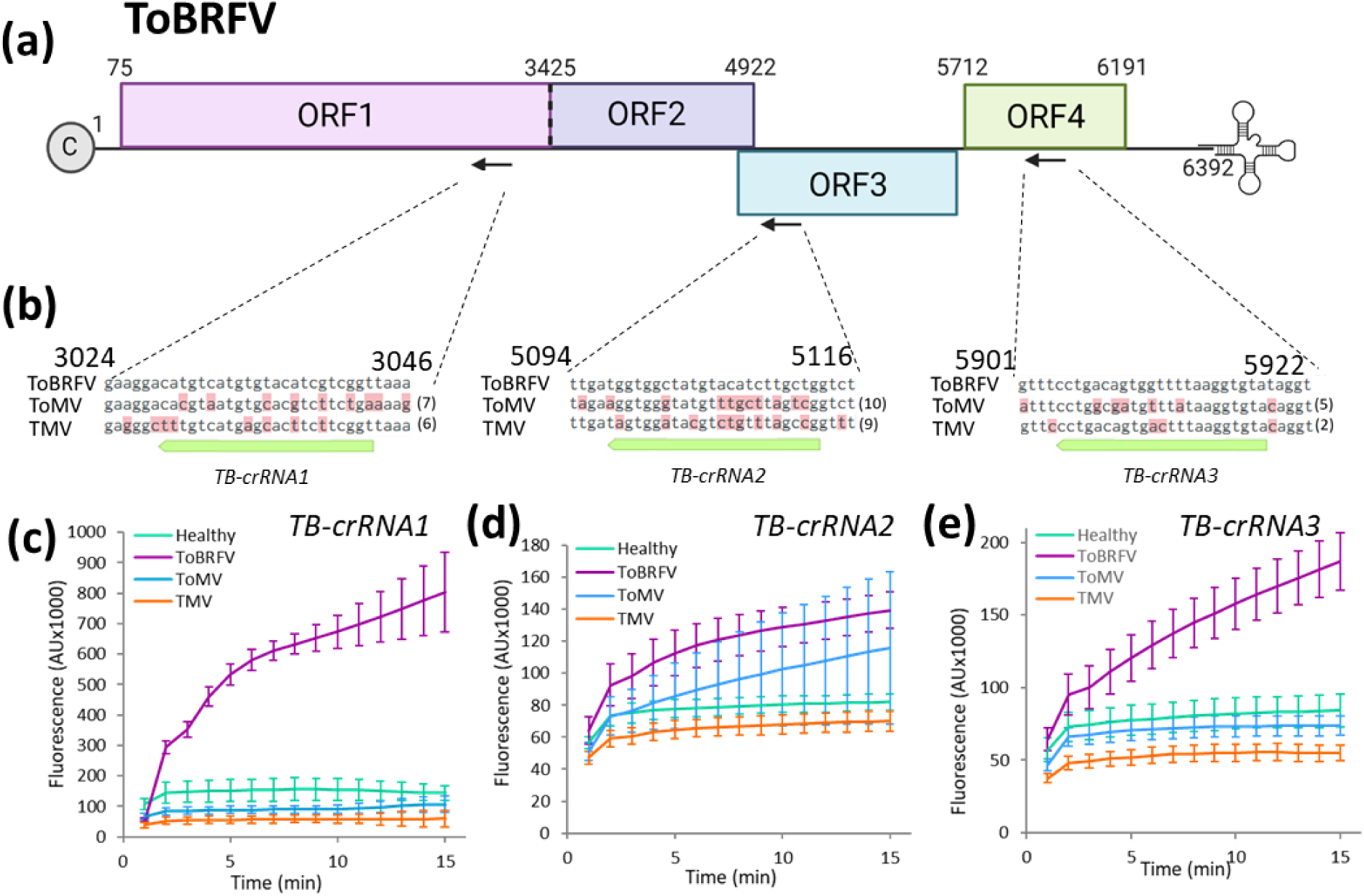
Analysis of different crRNAs for Cas13a-mediated direct detection of ToBRFV. **(a)** Schematic illustration of the ToBRFV genome, with the crRNA target sites. Three crRNAs were designed for ToBRFV detection: *TB-crRNA1* targeted ORF1/2 (nt 3024-3046), *TB-crRNA2* targeted ORF3 (nt 5090-5116) and *TB-crRNA3* targeted ORF 4 (nt 5901-5922). **(b)** Alignments of the ToBRFV crRNA target sites with the homologous sequences in ToMV and TMV. Mismatches are indicated in red and their number in parenthesis. **(c-e)** CRISPR/Cas13-based detection of ToBRFV using *TB-crRNA1* **(c)**, *TB-crRNA2* **(d)** and *TB-crRNA3* **(e)**. Detection was performed on RNA samples from healthy (green), ToBRFV- (purple), ToMV- (blue) and TMV- (orange) infected plants. Error bars indicate the standard error values of at least 4 biological replicates.

### CRISPR/Cas13a enables sensitive and early detection of ToBRFV

To examine the sensitivity of CRISPR/Cas13a-mediated ToBRFV detection, a series of dilutions of RNA samples from ToBRFV-infected plants with healthy plants was performed and subjected to Cas13a detection assays using *TB-crRNA1* (Figure 3a). Here, ToBRFV could be detected in 100-fold dilutions, suggesting that this technology could be used to detect one infected plant out of 100 via a sample pooling strategy. We further compared this technology with other detection methods. Detection of ToBRFV via Western-blot was efficient in the dilution of up to 5-fold of protein extracts from infected plants (Figure 3b; Supplementary figure S4), establishing that direct detection of ToBRFV using Cas13a is more sensitive than antibody-based methods. In contrast, RT-qPCR was able to detect ToBRFV in all tested dilutions (Figure 3b; Supplementary figure S4). In the 100-fold dilution, the Ct value of the RT-qPCR test was 17.5. Therefore, Cas13a-mediated detection is less sensitive than RT-qPCR.

**Figure 3.**
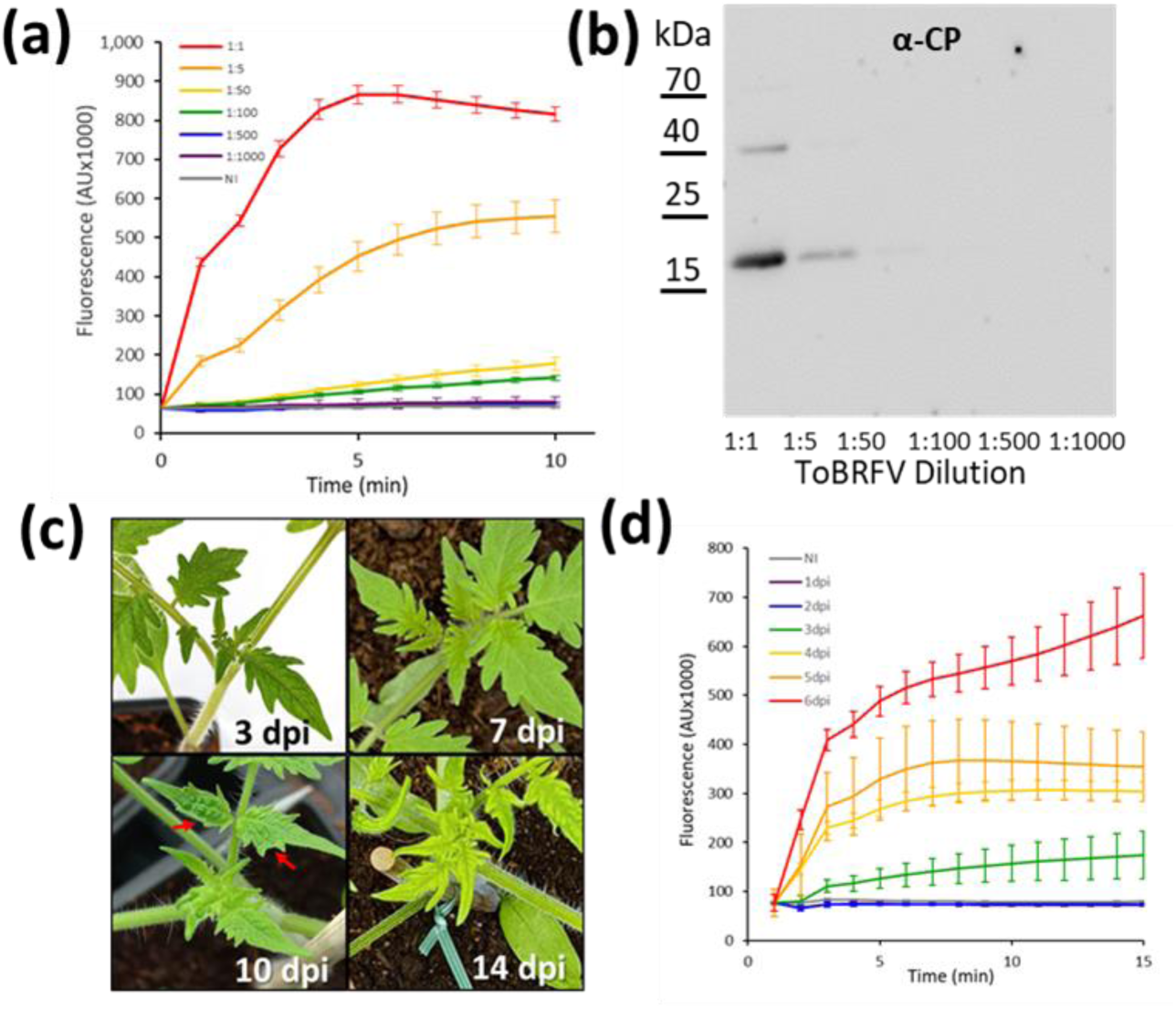
Sensitivity of ToBRFV CRISPR/Cas13a-mediated detection. **(a)** CRISPR/Cas13a detection of ToBRFV in serial dilutions of RNA samples from ToBRFV-infected plants diluted in RNA from healthy plants. Viral RNA can be detected in a 1:100 dilution. **(b)** Western blot analysis of coat protein (CP) in serial dilutions of protein samples from ToBRFV-infected plants. **(c)** Onset of ToBRFV symptoms during infection. First symptoms are observed at approximately 10 days post inoculation (dpi), indicated by red arrows. **(d)** CRISPR/Cas13a-mediated ToBRFV detection in systemic leaves at different dpi. Error bars indicate the standard error values of at least 5 biological replicates.

Timely detection of viral infection is required for the initiation of proper control measures. In ToBRFV-infected plants, symptoms start to develop approximately 10 days post-inoculation (dpi) (Figure 3c). To test if Cas13a is able to detect ToBRFV prior to the onset of symptoms, detection of ToBRFV in systemic leaves (distant from the site of infection) was examined at different time points after inoculation (Figure 3d). Strikingly, ToBRFV RNA could be detected in systemic leaves as early as 3 dpi (Figure 3d), long before the onset of symptoms. Collectively, these results establish that CRISPR/Cas13a-mediated detection is more sensitive than antibody-based methods, and could detect ToBRFV in up-to-100 fold dilutions and early after inoculation, prior to the development of disease symptoms.

### Detection of CGMMV in cucumber plants and TuMV in *Brassica napus* using CRISPR/Cas13

To determine the robustness of CRISPR/Cas13-mediated diagnosis, crRNAs were designed against two other agriculturally-significant viruses. The first was the cucurbit-infecting CGMMV. Since in ToMV and ToBRFV the most successful detection was for crRNAs targeting ORF1/2, two crRNAs (*CG-crRNA1* and *CG-crRNA2*) were designed to target the same region in CGMMV (Figure 4a). Detection by both crRNA was tested on RNA samples from healthy or CGMMV-infected cucumber plants (cv. Ilan). Indeed, *CG-crRNA1* and *CG-crRNA2* enabled effective detection of CGMMV (Figure 4b,c).

**Figure 4.**
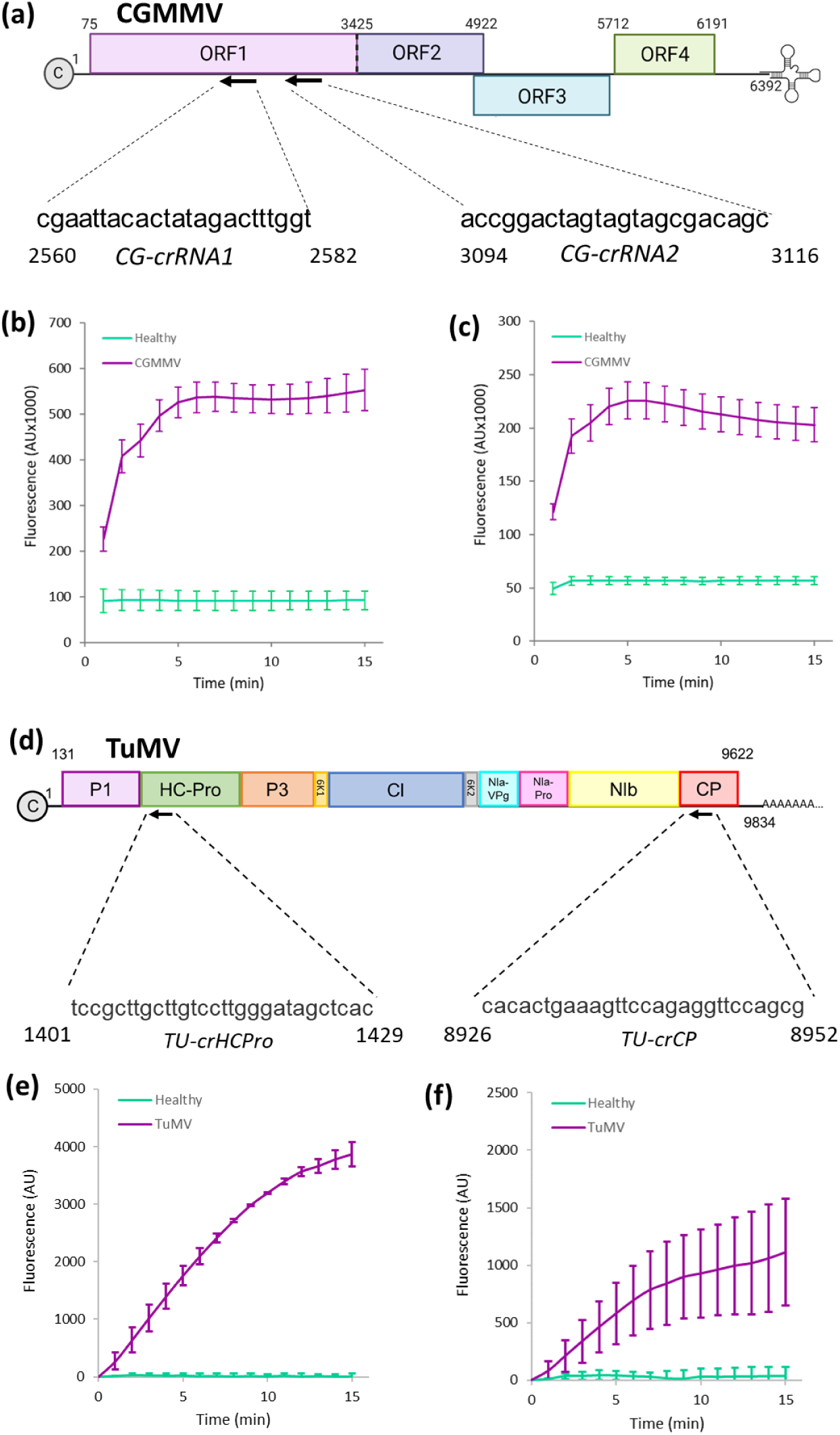
CIRSPR/Cas13a-mediated detection of CGMMV in cucumber and TuMV in rapeseed plants. **(a)** Schematic illustration of the CGMMV genome with the crRNA target sites and sequences. Two crRNAs were designed for CGMMV detection: *CG-crRNA1* (nt 2560-2582) and *CG-crRNA2* (nt 3094-3116). **(b-c)** CRISPR/Cas13-based detection of CGMMV using *CG-crRNA1* **(b)** and *CG-crRNA2* **(c)**. Detection was performed on RNA samples from healthy (green) and CGMMV-infected (purple plants. **(d)** Schematic illustration of the TuMV genome with crRNA sequences and target sites. crRNAs were designed according to Aman et al. (2018). **(e-f)** TuMV detection with crRNA targeting HC-Pro **(e)** and CP **(f)**. *TU-crHCPro* showed superior detection sensitivity and reproducibility compared to *TU*-*crCP*. Error bars indicate the standard error values of at least 4 biological replicates.

To test whether our CRISPR/Cas13a-mediated assay is applicable to other economically relevant viruses that do not belong to Tobamoviruses, we used rapeseed infected with TuMV and monitored the presence of viral RNA. The crRNAs were designed according to recently published studies on Cas13 mediated TuMV interference in plants (Aman *et al*., 2018), targeting RNA sequences within the HC-Pro (*TU-crHCPro*; nt 1401-1429) and CP (*TU-crCP*; nt 8926-8952) regions respectively (Figure 4d). In line with the findings of Aman et al. (2018), cRNA targeting the HC-Pro RNA region showed superior detection sensitivity and reproducibility in comparison to the crRNA targeting the CP RNA region (Figure 4e,f). These results suggest that Cas13a-mediated direct detection is applicable across families of viruses and crop species.

### A simplified approach for CRISPR/Cas13a-mediated ToBRFV detection in field samples

The robust ability of CRISPR/Cas13 to directly detect viruses from RNA extracts holds strong potential for on-site detection application. To simplify this process, we optimized a few steps in the detection protocol. First, a 5-minute protocol for crude RNA sampling was used (Marqués *et al*., 2022). To understand the sensitivity of this sampling method, a series of samples of ToBRFV-infected tomato leaf was performed and tested using the CRISPR/Cas13a protocol. With this extraction method, ToBRFV was detected in 1:5 dilutions with RNA from non-infected plants (Figure 5a).

**Figure 5.**
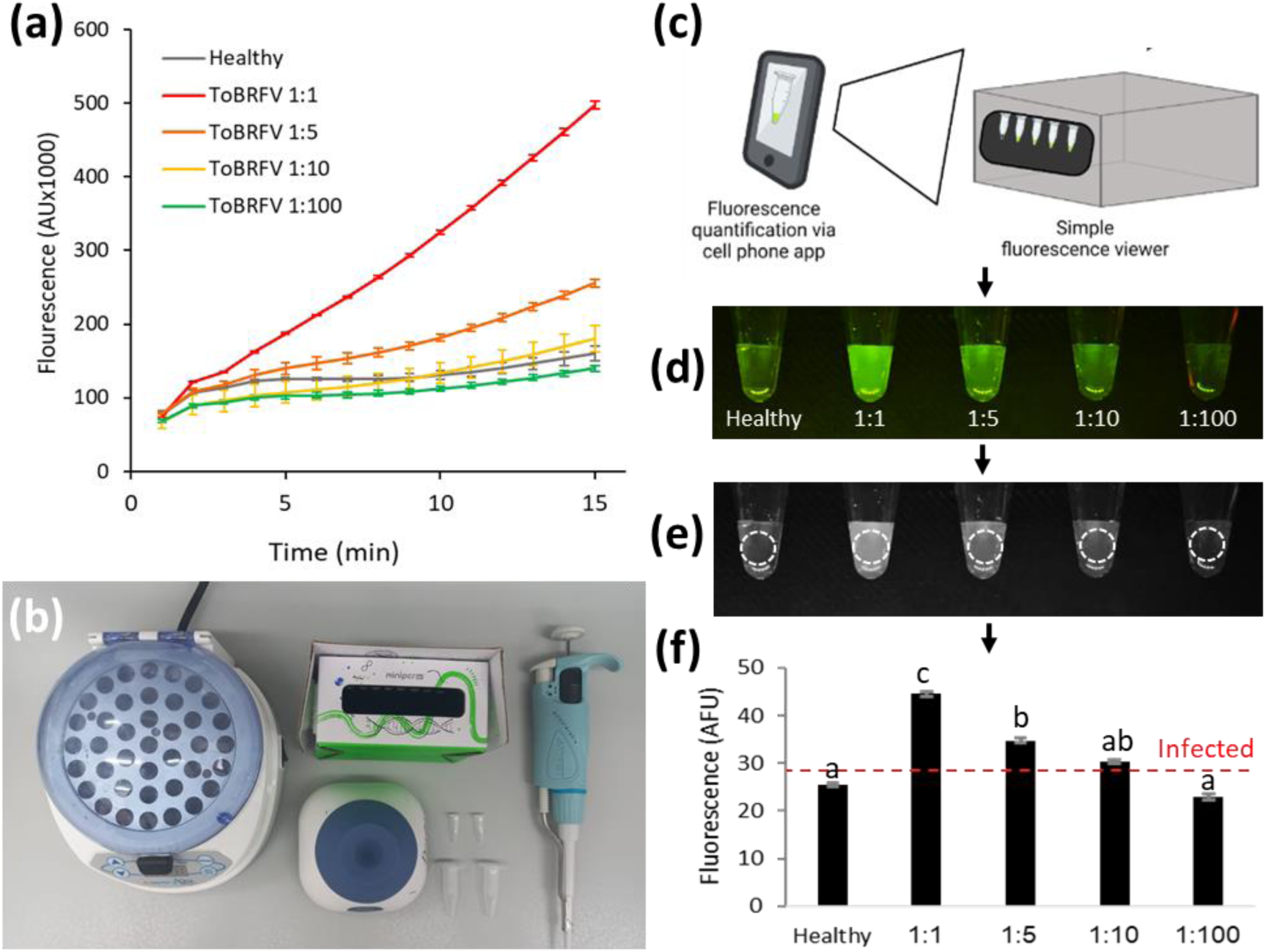
Development of a simplified approach for on-site Cas13a-based diagnostics. **(a)** Cas13a-mediated detection of ToBRFV in serial dilutions of crude RNA samples from ToBRFV-infected plants diluted in RNA from healthy plants. Viral RNA can be detected in a 1:5 dilution using this method. **(b)** Equipment required for on-site diagnostics includes dry bath, vortex, portable fluorescence viewer, a pipette and test tubes. **(c)** Schematic illustration of virus detection using a portable fluorescence viewer and a cellular phone camera. (d) Cellular phone camera image of serial dilutions of crude RNA from ToBRFV-infected plants. **(e)** Samples are converted to grayscale. **(f)** Fluorescence intensity is quantified using ImageJ. Quantification of serial dilutions shows detection at 1:5 dilution, similar to results achieved with a laboratory fluorescence reader. Error bars indicate the standard error values of at least 4 biological replicates. Different letters indicate statistical significance in Tukey’s-HSD test.

Next, we utilized a simple, affordable ($35) portable and battery-operated fluorescence viewer to detect the Cas13 signal (Figure 5b). The fluorescent tubes were then imaged using a cellular phone camera (Figure 5c). The fluorescent response was captured (Figure 5d), transformed to grayscale (Figure 5e), and quantified using the ImageJ software (Figure 5f). Strikingly, this approach yielded sensitivity similar to that obtained with laboratory equipment, detecting ToBRFV in up-to 5-fold dilutions of RNA from infected plants. This direct fluorescence detection and quantification method was also confirmed for CGMMV detection in cucumber and TuMV detection in rapeseed plants (Supplementary figure S5), suggesting it can be applied for different virus-crop pathosystems.

To test the validity of this approach for ToBRFV diagnosis in field samples, a blind test was performed. In March 2023, an outbreak of an unknown virus occurred in Yated village, southern Israel. The symptoms were consistent with tobamovirus infection (Figure 6a). RNA was sampled from leaves of infected plants and subjected to CRISPR/Cas13-mediated direct identification of ToBRFV or ToMV using *TB-crRNA1* or *TM-crRNA*, respectively. These tests identified the virus as ToBRFV, and not ToMV (Figure 6b,c). The presence of ToBRFV was further confirmed by RT-PCR followed by sequencing analysis (Supplementary figure S6). This result demonstrated the high applicability of Cas13a-mediated detection for rapid, direct, and specific on-site detection.

**Figure 6.**
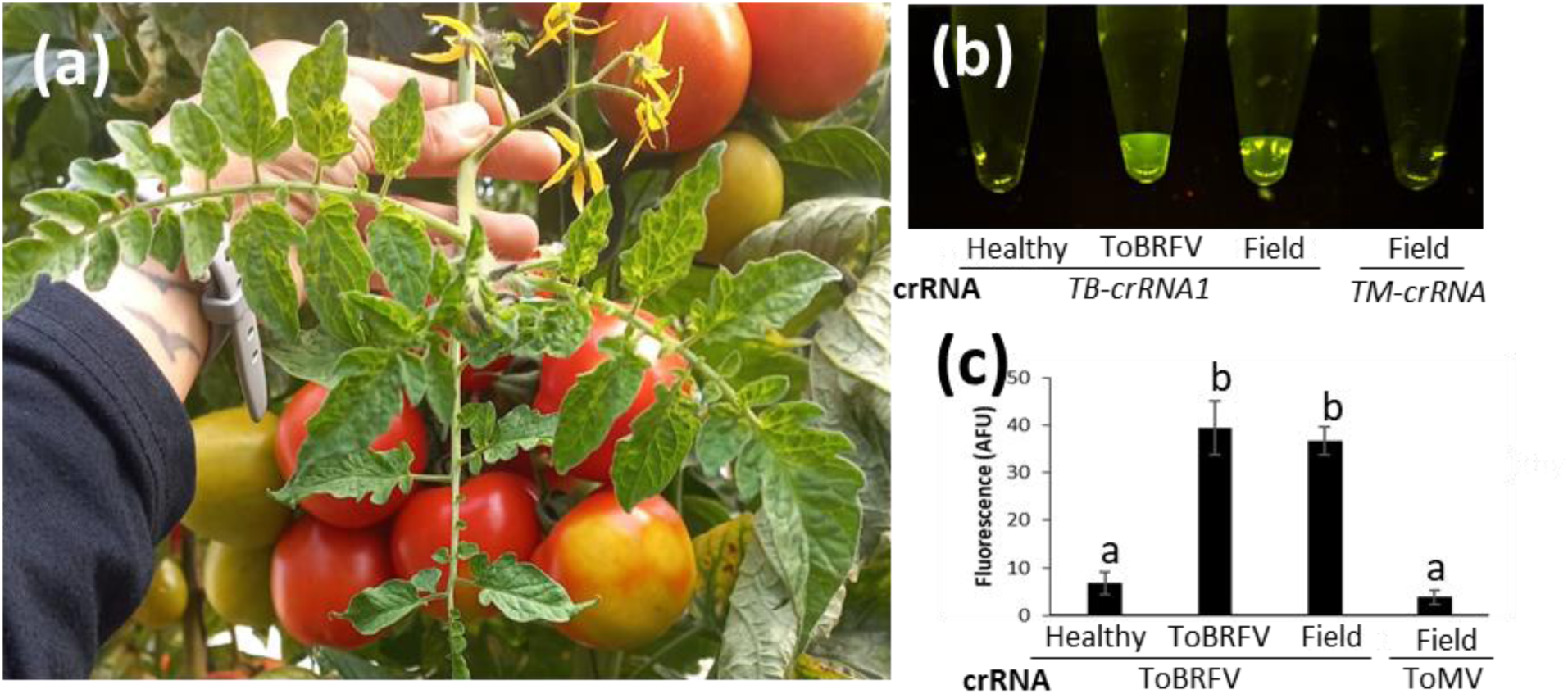
Cas13-based on-site diagnostics. **(a)** Image from a tomato virus outbreak in Yated village, southern Israel. **(b)** Cas13-based detection of ToBRFV in the suspected filed sample. **(c)** Quantification of the fluorescent signal. Different letters indicate statistical significance in Tukey’s-HSD test.

## Discussion

Proper management of crop viral diseases requires rapid, specific, and sensitive identification of the causative agent. The emergence of new and resistance-breaking viruses requires new tools that will enable rapid detection as a first step toward mitigating their dispersal. Here, we present a simple protocol for direct identification of viruses using Cas13a (Figure 1). Our results suggest that Cas13a could be a useful tool for rapid, amplification-free, and sequence-specific detection of several viruses in various crop species. Rapid detection of ToBRFV using Cas13a specifically distinguished between this emerging virus and the closely related viruses TMV and ToMV (Figure 2). Cas13a-mediated detection was sensitive, and allowed the detection of ToBRFV in 100-fold dilution of RNA from infected plants and at an early point after inoculation, a week prior to the onset of symptoms (Figure 3). By changing the crRNA sequence, this system could be adapted to the identification of various viruses in different crop species, including CGMMV in cucumber and TuMV in *B. napus* (Figure 4). In addition, we show that Cas13a-mediated detection can be conducted using a $35, simple, portable fluorescence viewer (Figure 5) and that it can be used to detect ToBRFV in field samples (Figure 6). These results establish Cas13a as a useful method for rapid, sensitive, and sequence-specific detection of plant RNA viruses.

Previous studies have demonstrated the ability of Cas13a for virus identification. The pioneering development of Cas13a detection for various human disease-causing viruses demonstrated the vast potential of this technology (Freije *et al*., 2019; Metsky *et al*., 2020; Myhrvold *et al*., 2018). It is worth noting that all Cas13a detection methods of human viruses included an amplification step, and did not allow direct detection from extracted RNA. In plants, expression of CRISPR-Cas13 was demonstrated to confer resistance to various RNA viruses (Mahas *et al*., 2019) and allowed the detection of plant genes (Abudayyeh *et al*., 2019). However, Cas13-mediated detection of plant viruses was demonstrated in only a few cases. Cas13a-mediated detection was successfully applied for tomato spotted wilt virus in its tomato host and thrips vector *Frankliniella occidentalis*, however, this identification also relied on amplification of the viral RNA (Zhang *et al*., 2021). Recently, direct identification of RNA viruses in infected plants was shown in the model plant *Nicotiana benthamiana* (Marqués *et al*., 2022), however, it was not yet demonstrated in crop systems and still depended on laboratory fluorescence detectors. In tomato, direct identification of RNA viruses was recently shown using CRISPR-based amplification-free digital RNA detection (SATORI), however, this system also required specialized equipment for detection (Ueda *et al*., 2023). Here, we demonstrate a rapid, amplification-free approach, which does not require any specialized equipment such as PCR or laboratory fluorescence reader. In addition, we provide evidence for the robustness of this system by applying it to multiple crop species and viruses of different families. We propose that this method could serve as a platform for the rapid, accurate, and sensitive diagnostics of various viruses in the field or in control points for international commerce of fruit, seeds, and propagation material.

Our on-site identification protocol is parallel in its applicability and detection time to existing on-site rapid detection methods based on antibodies. However, there are several important advantages to Cas13a-based detection. First, it allows sequence-specific detection, as shown by the distinction of ToBRFV from closely-related tobamoviruses. This advantage can be further used for the distinction between different strains, isolated and for the detection of resistance-breaking mutations (Shymanovich *et al*., 2024). Another important advantage is the modularity of Cas13-based detection. The production and optimization of antibody-based protocols is a challenging and lengthy process. The generation of less-specific polyclonal antibodies often involves the use of experimental animals, lasts several months, and requires extensive quality control. Monoclonal antibodies are more specific (Bernabé-Orts *et al*., 2021), but their production is more complex, challenging, and time-consuming. Using the Cas13a method, any crRNA can easily be generated against any sequence. In the future, this technology may be able to replace antibody-based methods for on-site detection of plant RNA viruses.

## Experimental procedures

### Plant material and virus inoculation

Tomato (*Solanum lycopersicum* cv. Moneymaker) (LA2706), cucumber (*Cucumis sativus* cv. Ilan) and *Nicotiana benthamiana* plants were grown in soil in a light- and temperature-controlled chamber at 25 °C with a 16 h light/8 h dark regime. Rapeseed plants (Brassica napus cv. Drakkar) were grown in soil in a greenhouse at 20 °C. For the ToBRFV time-course experiments, tomato plants were grown in a greenhouse at 25 °C. Ten-day-old tomato plants were used for mechanical inoculation of ToBRFV or ToMV. For ToBRFV infection, we used the Israeli isolate of ToBRFV, ToBRFV-IL (KX619418.1). ToMV was derived from the infectious clone pTLW3 (Hak *et al*., 2023). Plants were rubbed with a phosphate buffer solution (0.01 M, pH 7.0) supplemented with carborundum powder and crushed leaves from ToMV- or ToBRFV-infected tomato plants. For TMV infection, four-week-old *N. benthamiana* plants were agro-infiltrated with the vector pJL24 that harbors the TMV genome and a GFP reporter (TMV-GFP) (Lindbo, 2007). For CGMMV, one-week-old cucumber plants were agro-infiltrated with a CGMMV infectious clone plasmid (Liu *et al*., 2017). TuMV (isolate UK1) infectious clone plasmid (pCB-TuMV-GFP, (Garcia-Ruiz *et al*., 2012) was agro-infiltrated into three-week-old *Brassica napus* plants.

### Agroinfiltration

*Agrobacterium tumefaciens* EHA105 for *N. benthamiana*, or GV3101 for cucumber, harboring a binary vector were grown overnight in a 28°C shaker. Cell cultures were harvested and resuspended in morpholineethane sulfonic acid (MES) buffer (10 mM MgCl2, 10 mM MES, 150 μM acetosyringone, pH 5.6) to an OD600 of 0.3 and were infiltrated, using a needleless syringe, into the abaxial side of the fourth or fifth leaf of *N. benthamiana* plants or cotyledons of cucumber seedlings.

### RNA extraction

Total plant RNA was extracted from 50-100 mg of systemic leaf tissue of virus-infected plants using Total RNA Mini Kit (Geneaid, RPD050, Taiwan R.O.C). Eluted RNA was diluted to a concentration of 100 ng/μl. Crude RNA sampling was done as previously described (Marqués *et al*., 2022). Briefly, leaf segments of 50 mg were placed in 300 μL of lysis buffer (15% polyethylene glycol 4000, 20 mM NaOH), incubated for 5 min at room temperature, vortexed and placed on ice.

### Cas13a protein expression

*L. buccalis* Cas13a (LbuCas13a) was produced and purified as recently published (Fozouni *et al*., 2020) with minor modifications. In brief, *E.coli* Rosetta 2 (DE3) (Merck, Darmstadt, Germany) cells harboring the plasmid p2CT-His-MBP-Lbu_C2c2_WT (Addgene No. 83482) were cultivated with 800 ml autoinduction medium (Studier, 2005). After an initial 3-hour incubation at 37 °C with shaking at 190 rpm, cultures were further grown overnight at 24 °C. Cells were harvested by centrifugation at 5700 x g for 20 minutes at 4°C and stored at −20 °C until further use. Pellets were resuspended in 70 ml ice-cold lysis buffer (50 mM Tris-HCl pH 7.0, 500 mM NaCl, 5 % (v/v) glycerol, 1 mM TCEP, 1 mM AEBSF) and 1 protease inhibitor tablet (Merck). Lysis was done at 4 °C with the addition of 1 mg/ml lysozyme and further incubation at 4 °C for 30 minutes, followed by sonication using a Branson sonifier 250 (8 repeated cycles 30 seconds on, 20 seconds off, 50% duty cycle, 50 % energy output). After subsequent centrifugation at 30000 x g for 30 minutes at 4°C, supernatants were filtered through a 0.45 µm syringe filter. The filtrate was loaded onto a 5 ml HisTrap FF column (Cytiva, Uppsala, Sweden) and washed over 10 column volumes (CV) with lysis buffer containing 1 M NaCl followed by a gradient elution with buffer B (50 mM Tris-HCl pH 7.0, 500 mM NaCl, 5 % (v/v) glycerol, 1 mM TCEP, 1 M imidazole) over 15 CV. Fractions containing 6xHis-MBP-LbuCas13a were pooled and dialyzed overnight at 4 °C against 50 mM Tris-HCl pH 7.0, 250 mM KCl, 5 % (v/v) glycerol, 1 mM TCEP. TEV protease (Cytiva) was added for tag removal. Cleaved LbuCas13a was further purified via size exclusion chromatography using a HiLoad Superdex 200 pg column (Cytiva) equilibrated with 20 mM HEPES pH 7.0, 200 mM KCl, 10 % (v/v) glycerol and 1 mM TCEP. Fractions were pooled and concentrated to a final concentration of 2 mg/ml using a centrifugal filter device (Sartorius, Göttingen, Germany) with a molecular weight cut-off of 50 kDa.

### Fluorescent Cas13 nuclease assay

Twenty-three bp-long specific crRNAs were designed using the Cas13 design tool https://cas13design.nygenome.org/ fused to the 3’ end of the LbuCas13a stem (5’ GACCACCCCAAAAAUGAAGGGGACUAAAAC 3’), as in (Fozouni *et al*., 2020). Guide ToBRFV crRNA1 and ToMV crRNA were designed with the same sequence as for Cas12a (Alon et al. 2021). ToBRFV guides were chosen to have at least 5 mismatches when compared to ToMV (see oligo table). RNA oligonucleotides were synthesized by Integrated DNA Technologies (IDT, Coralville, USA).

RNPs were assembled by individually mixing 1 μM of each crRNA oligo with 1 μM LbuCas13a and incubating for 30 minutes at room temperature. Cleavage reaction of a total volume of 10 μL was done using 300 ng total RNA (or 3 μL of crude RNA), 100 nM RNPs, and 200 μM RNaseAlert® substrate (IDT, Coralville, USA) as reporter in cleavage buffer (20 mM Hepes pH 6.8, 50 mM KCl, 5 mM MgCl2, 5% Glycerol). The reaction was incubated at 37°C for up to 30 minutes, and FAM reading was taken every minute in a QuantStudio3 (Thermo) instrument. For on-site detection, fluorescence was viewed using the P51™ Molecular Fluorescence Viewer from MiniPCR Bio (Cambridge, MA, USA), and fluorescence was quantified using ImageJ (https://imagej.net/ij/).

### Reverse transcription quantitative PCR (RT-qPCR)

Total RNA from ToBRFV-infected plants (500 ng) served as a template for cDNA synthesis using the qPCRBIO cDNA Synthesis Kit, according to the manufacturer’s protocol (PCRBIO, cat. no. PB30.11). qRT-PCR was performed using 2×SyGreen Mix (PCRBIO) In QuantStudio3 instrument (Thermo). Virus levels were calculated as the expression ratio of ToBRFV-CP and the reference gene *TIP1* (see table for primer sequences).

### Evaluation of ToBRFV levels by Western blot

For detection of ToBRFV in tomato plants, the first developing leaf (1 to 3 cm) and apex were collected and ground, while frozen, in a microcentrifuge tube. Laemmli buffer (40 μl of 3×) (100 mM Tris, 2% SDS, 20% glycerol, 4% β-mercaptoethanol, pH 6.8) was added and mixed into the sample, followed by centrifugation for 10 min and boiling of the supernatant for 5 min. Samples were then run on 12% SDS-PAGE acrylamide gels, transferred to nitrocellulose membranes (Protran), and blocked with 3% skimmed milk in Tris buffer saline-Tween. CP of the virus was detected by rabbit anti-tobamovirus CP (1:20,000), courtesy of Dr. A. Dombrovsky, ARO, and anti-rabbit horseradish peroxidase (1:20,000) (Jackson immunoresearch). Chemiluminescence was observed using Elistar Supernova as substrate (Cyanagen), and images of protein bands were acquired and quantified using the Alliance Q9 software (UVITEC).

## Accession numbers

ToBRFV-IL KX619418.1; CGMMV NC_001801.1; TuMV NC_002509.2;

## Acknowledgements

This work was supported by the Israeli Ministry of Agriculture and Rural Development (GA No. 20-02-0187) and the National Center for Genome Editing Applications and Technologies in Agriculture (GA No. 20-01-0209). This project has also received funding from the European Research Council (ERC) under the European Union’s Horizon 2020 research and innovation programme (GA No. 810131) and from the Deutsche Forschungsgemeinschaft (DFG; GA No. 433194101, Research Unit 5116).

## Author contributions

H.H., S.O., G.P., J.K. and Z.S. conceptualized and designed the experiments; H.H., S.O. and A.R. performed the experiments; S.I.G. detected and provided field samples; H.H., S.O., A.R., G.P., J. K. and Z.S. analyzed the data; H.H., S.O., G.P., J.K. and Z.S. wrote the manuscript;

