## Supplementary figure for "Rapid, direct, and sequence-specific identification of RNA viruses in various crop plants using CRISPR/Cas13a"

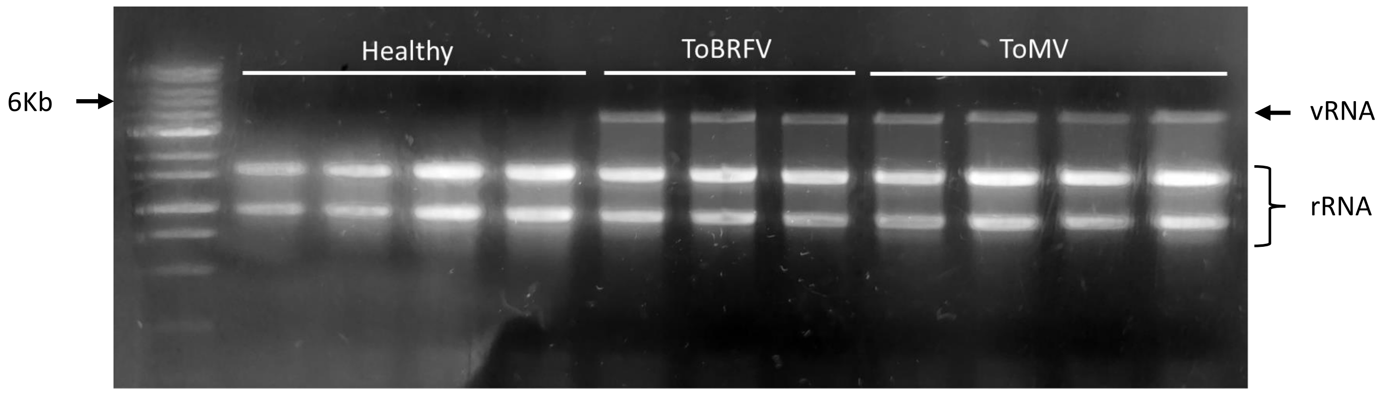


**Supplementary figure S1. Accumulation of tobamovirus viral RNA in leaf tissue.** Agarose gel electrophoresis of RNA samples from healthy, ToBRFV- and ToMV- infected leaf tissue. The viral genomic RNA (vRNA) can be easily observed at ~6.4kB.

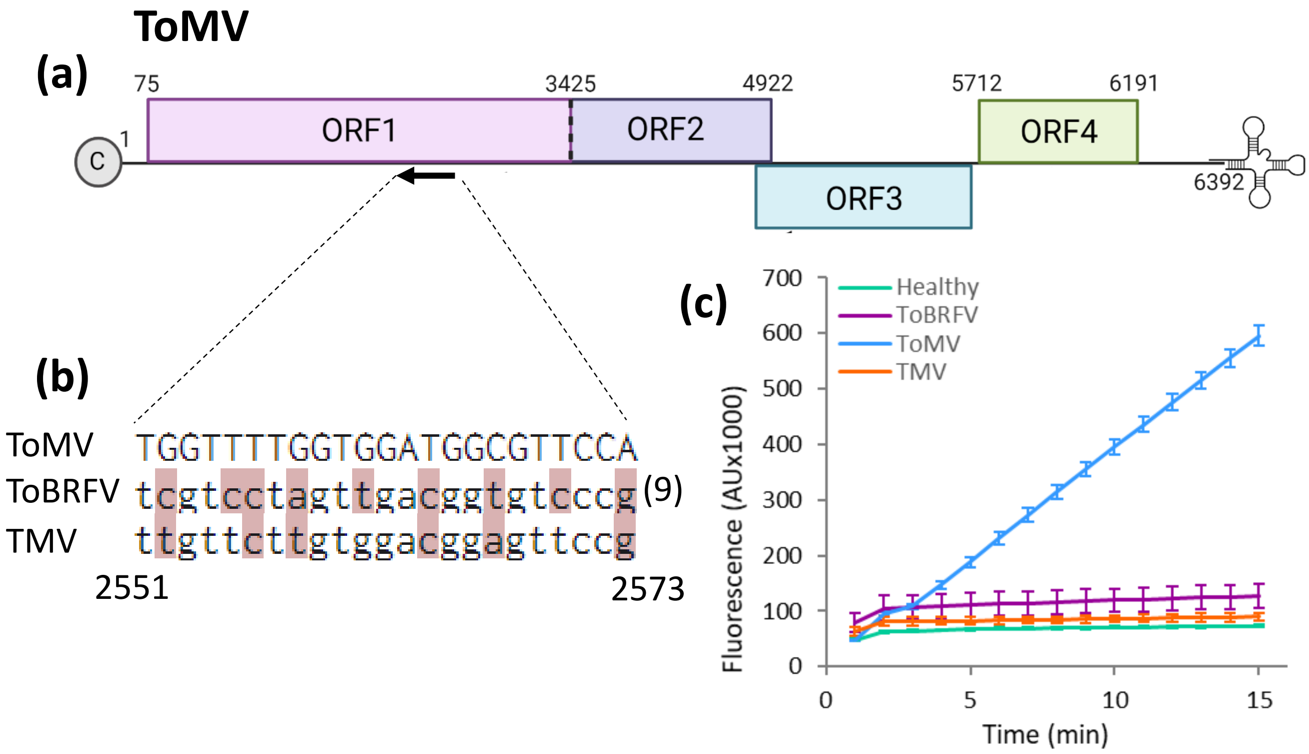


**Supplementary figure S2. Cas13a-mediated direct detection of ToMV.** (a) Schematic illustration of the ToMV genome, with the crRNA target site. Three crRNAs were designed for ToBRFV detection: TB-crRNA1 targeted ORF1/2 (nt 2551-2573. (b) Alignment of the ToMV crRNA target sites with the homologous sequences in ToMV and TMV. Mismatches are indicated in red and their number in parenthesis. (c-e) CRISPR/Cas13-based detection of ToBRFV using crRNA 1 (c), crRNA 2 (d) and crRNA 3 (e). Detection was performed on RNA samples from healthy (green), ToBRFV- (purple), ToMV- (blue) and TMV- (orange) infected plants. Error bars indicate the standard error values of at least 4 biological replicates.

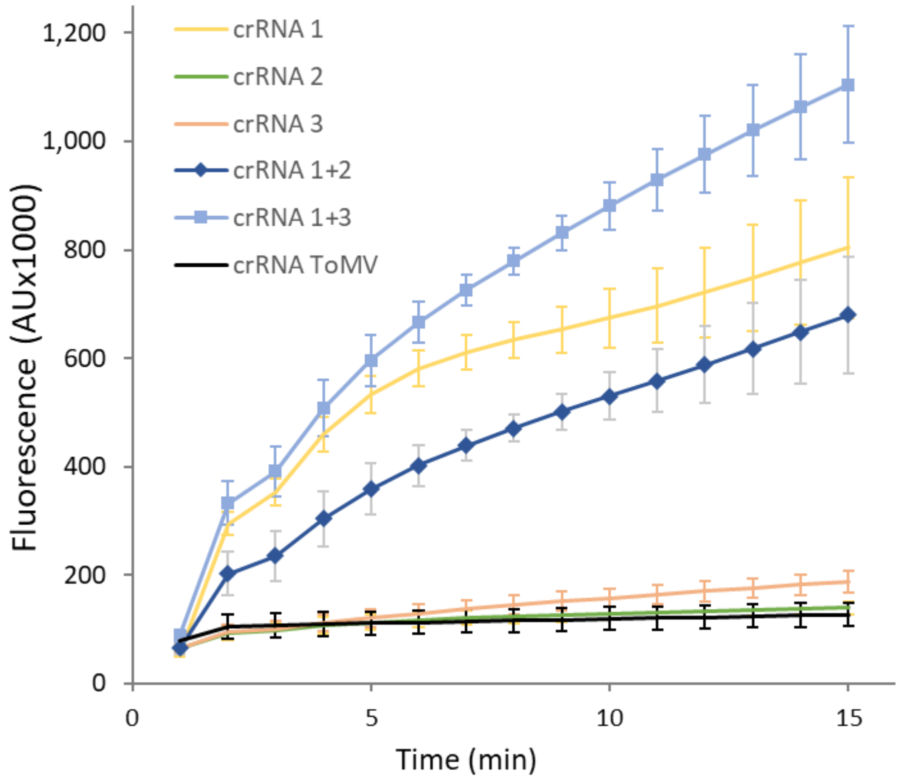


**Supplementary figure S3. Combination of different crRNAs enhances the Cas13-emitted ToBRFV signal.** CRISPR/Cas13-based detection of ToBRFV using TB-crRNA 1 (yellow), TB-crRNA2 (green), TB-crRNA3 (orange), a combination of TB-crRNA1+2 (dark blue), a combination of TB-crRNA1+3 (light blue) and TM-crRNA (black). Detection was performed on RNA samples from ToBRFV-infected plants. Error bars indicate the standard error values of at least 4 biological replicates.

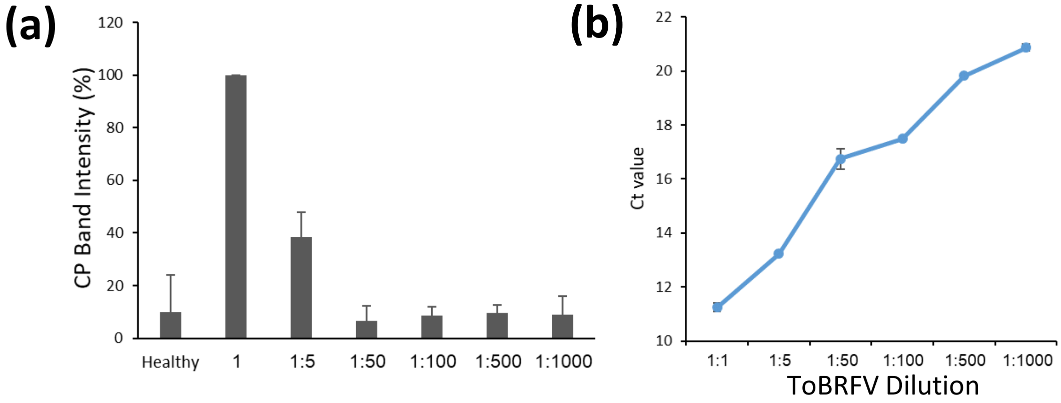


**Supplementary figure S4. Detection of a serial dilution of ToBRFV using RT-qPCR. (a)** Quantitative analysis of Western blot band intensity in a serial dilution of ToBRFV-infected plant protein samples. **(b)** Analysis of RT-qPCR assay Ct values in serial dilutions of samples from ToBRFV-infected plants with RNA from healthy plants.

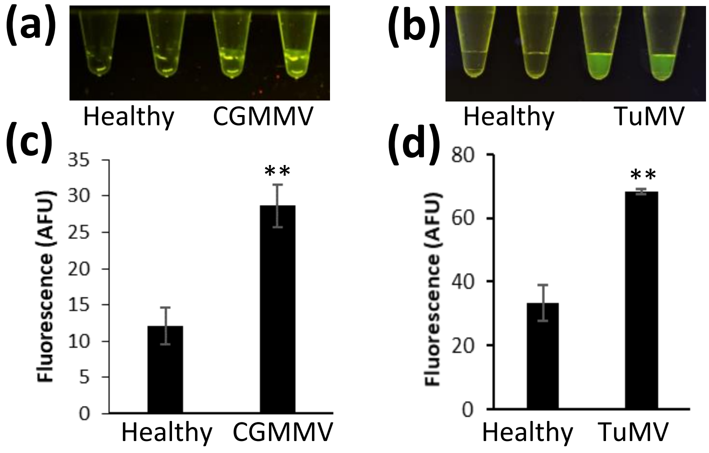


**Supplementary figure S5. Visual detection of CGMMV and TuMV. (a-b)** visual Cas13a-based detection of CGMMV in RNA samples from cucumber plants **(a)** and TuMV from rapeseed plants **(b)** using a fluorescence viewer. **(c-d)** Quantification of fluorescence in, CGMMV- **(c)** and TuMV-infected plant RNA samples **(d)** using ImageJ. Error bars indicate the standard error values of at least 3 biological replicates.

(a)

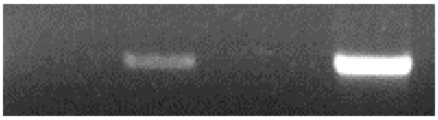


(b)

```
ToBRFV-IL       -------------ATGGCTCTTGTTAAGGGTAAAGTCAATATTAATGAGTTCATAGACTT
Field_sample    TTTGAAGACAAAAATGGCTCTTGTTAAGGGTAAAGTCAATATTAATGAGTTCATAGACTT
                             ***********************************************

ToBRFV-IL       GTCAAAATCAGAAAAATTTCTTCCGTCTATGTTCACACCTGTTAAGAGTGTCATGATCTC
Field_sample    GTCAAAATCAGAAAAATTTCTTCCGTCTATGTTCACACCTGTTAAGAGTGTCATGATCTC
                ************************************************************

ToBRFV-IL       CAAGGTTGATAAGATATTGGTTCATGAAGATGAATCTTTGTCCGAAGTCAATTTACTCAA
Field_sample    CAAGGTTGATAAGATATTGGTTCATGAAGATGAATCTTTGTCCGAAGTCAATTTACTCAA
                ************************************************************

ToBRFV-IL       AGGTGTAAAACTCATTGATGGTGGCTATGTACATCTTGCTGGTCTTGTGGTGACAGGTGA
Field_sample    AGGTGTAAAACTCATTGATGGTGGCTATGTACATCTTGCTGGTCTTGTGGTGACAGGTGA
                ************************************************************

ToBRFV-IL       ATGGAATTTGCCAGATAATTGTCGTGGTGGTGTCAGTGTCTGTTTGGTCGATAAGAGAAT
Field_sample    ATGGAATTTGCCAGATAATTGTCGTGGTGGTGTCAGTGTCTGTTTGGTCGATAAGAGAAT
                ************************************************************

ToBRFV-IL       GGAGAGAGCGGACGAGGCAACTCTTGCTTCATACTATACCGCAGCGGCTAAGAAAAGGTT
Field_sample    GGAGAGAGCGGACGAGGCAACTCTTGCTTCATACTATACCGCAGCGGCTAAGAAAAGGTT
                ************************************************************

ToBRFV-IL       TCAGTTCAAAGTCGTTCCAAATTACAACATCACTACCAAGGACGCAGAAAAGGCAGTTTG
Field_sample    TCAGTTCAAAGTCGTTCCAAATTACAACATCACTACCAAGGACGCAGAAAAGGCAGTTTG
                ************************************************************

ToBRFV-IL       GCAAGTACTAGTTAATATTAGAAATGTTAAAATTGCTGCGGGTTACTGTCCGCTGTCATT
Field_sample    GCAAGTACTAGTTAATATTAGAAATGTTAAAATTGCTGCGGGTTACTGTCCGCTGTCATT
                ************************************************************

ToBRFV-IL       AGAATTTGTGTCAGTGTGTATTGTTTATAAAAATATTATAAAACTCGGTTTGAGAGAGAA
Field_sample    AGAATTTGTGTCAGTGTGTATTGTTTATAAAAATATTATAAAACTCGGTTTGAGAGAGAA
                ************************************************************

ToBRFV-IL       AATTACGAGCGTCACGGATGGAGGGCCCATGGAACTATCAGAAGAAGTTGTTGATGAGTT
Field_sample    AATTACGAGCGTCACGGATGGAGGGCCCATGGAACTATCAGAAGAAGTTGTTGATGAGTT
                ************************************************************

ToBRFV-IL       CATGGAAGAAGTCCCGATGTCTGTAAGGCTTGCAAAATTTCGTTCGAAGACCGGAAAAAA
Field_sample    CATGGAAGAAGTCCCGATGTCTGTAAGGCTTGCAAAATTTCGTTCGAAGACCGGAAAAAA
                ************************************************************

ToBRFV-IL       GTTTAGTAGTAAAAGTGAGAATAATAGTGGTAATAATAGGCCGAAACCAAACAAAAACCA
Field_sample    GTTTAGTAGTAAAAGTGAGAATAATAGTGGTAATAATAGGCCGAAACCAAACAAAAACCA
                ************************************************************

ToBRFV-IL       AAGGAAGGAAAAGGGTTTAAAAGTTAGGGTTGAGAAGGATAATTTAATTGATAATGAATT
Field_sample    AAGGAAGGAAAAGGGTTTAAAAGTTAGGGTTGAGAAGGATAATTTAATTGATAATGAAT-
                ***********************************************************

ToBRFV-IL       GGAGACTTACGTCGCCGATTCAGATTCGTATTAA
Field_sample    GGAGACTT--------------------------
                ********
```

**Supplementary figure S6. Detection of ToBRFV in field samples.** (a) RT-PCR analysis of the field sample using ToBRFV-specific primers. Healthy, ToBRFV- and ToMV- infected plants served as controls. (b) Alignment of the sequencing result of the ToBRFV obtained from the field sample with ToBRFV-IL (KX619418.1).
